# Testing for reciprocal trait influence in plant-frugivore interactions using generalized joint attribute modeling

**DOI:** 10.1101/2024.01.17.576112

**Authors:** Laurel R. Yohe, Leith B. Miller, Zofia A. Kaliszewska, Susan R. Whitehead, Sharlene E. Santana, Liliana M. Dávalos

**Author notes:** **E-mail Addresses:** LRY;, LBM, ZAK, SRW, SES. **Statement of Authorship:** LBM, ZAK, SRW and SES collected primary data, and LRY and LMD designed and performed the analyses. All authors contributed to the discussion of the results and the writing of the manuscript. **Data Accessibility Statement:** All data, scripts, and results will be deposited onto Dryad. **Corresponding Authors:** Laurel Yohe 9331 Robert D. Snyder Rd. Charlotte, NC 28223 Liliana M. Dávalos 650 Life Sciences Stony Brook, NY 11794.

## Abstract

Under an adaptive hypothesis, the reciprocal influence between mutualistic plants and frugivores is expected to result in dispersal syndromes comprising both frugivore and plant traits that structure fruit consumption. Tests of this adaptive hypothesis, however, focus on traits of either fruits or frugivores but not both and often ignore within-species variation. To overcome these limitations, we analyze traits for the mutualistic ecological network comprising *Carollia* bats that feed on and disperse *Piper* seeds. For these analyses, we use generalized joint attribute modeling (GJAM), a Bayesian modeling approach that simultaneously accounts for multiple sources of variance across trait types. In support of the adaptive hypothesis and indicating niche partitioning among *Carollia* bats, we find differential consumption of a suite of *Piper* species influenced by bat traits such as body size; however, *Piper* morphological traits had no effect on bat consumption. Slow evolutionary rates, dispersal by other vertebrates, and unexamined fruit traits, such as *Piper* chemical bouquets, may explain the lack of association between bat *Piper* consumption and fruit morphological traits. We have identified a potential asymmetric influence of frugivore traits on plant-frugivore interactions, providing a template for future trait analyses of plant-animal networks.

## INTRODUCTION

A reciprocal influence between frugivore and fruit traits is often expected in ecological interactions comprising seed dispersers and plant mutualists (Janson 1983). But plant and animal populations may be generalists and within-population variation can obscure how organismal traits influence such interactions. Further, across rich ecological networks such as those in the Neotropics, an adaptive hypothesis for traits linking plant and animal species may be unwarranted. Instead, ecological fitting, whereby fruit-frugivore interactions emerge through the maintenance of ancestral traits in a new environment, could explain contemporary interactions without the need to invoke adaptation (Janzen 1985). Nevertheless, there is support for animals shaping fruit traits, or the dispersal syndrome hypothesis, in the form of fruit or seed size, hardness, color, scent and chemical profile matching frugivore preferences (Valenta and Nevo 2020). Conversely, vertebrate adaptations to frugivory are well supported, including sensory, digestive, and even excretory traits (Herrera 1984; Schondube et al. 2001; Saldaña-Vázquez et al. 2013; Wang et al. 2020; Yohe et al. 2021), which contribute to structuring mutualistic networks.

Testing contrasting adaptive and non-adaptive hypotheses is further complicated by the multiple scales at which interactions and traits are measured. While the selective influence of frugivory on plants has been examined through seed dispersal and recruitment analyses (Norconk et al. 1998; Nathan and Muller-Landau 2000; Howe and Miriti 2004), its effects on frugivore traits have been analyzed at scales that range from individuals to clades (Pratt and Stiles 1985; Stevenson et al. 2000; Burns 2004). Important gaps emerge from this variation in scales. In contrast to plant-pollinator interactions, the reciprocal influence of frugivory on plants and frugivores has seldom been tested simultaneously, and some studies overlook within-species variation. To date, research on the evolutionary consequences of fruit-frugivore interactions has primarily focused on traits, such as vertebrate color vision and fruit color indicating ripeness, that explain the foraging behavior of birds and diurnal mammals (Osorio et al. 2004; Schaefer et al. 2007). In contrast, the influence of fruit traits on nocturnal frugivores (e.g., bats) and vice versa is largely unknown (Luft et al. 2003; Hodgkison et al. 2013; but see Thies and Kalko 2004), even though bats constitute a large percentage of seed dispersers in tropical ecosystems (Fleming and Heithaus 1981; Muscarella and Fleming 2007; Fleming and John Kress 2011).

To test these broad hypotheses, neotropical *Piper* plants (Piperales: Piperaceae) and *Carollia* bats (Fig. 1; Chiroptera: Phyllostomidae) are an ideal mutualistic system whose ecology has been well documented. *Piper* are both diverse and abundant in tropical ecosystems worldwide (Gentry 1988) and provide a constant supply of ripe fruit throughout the year through continuous or staggered fruiting patterns among sympatric species (Thies and Kalko 2004). Many neotropical *Piper* species depend on *Carollia* for seed dispersal (Dyer and Palmer 2004), and *Piper* fruits dominate these bats’ diets throughout the year and across their range (Fleming 1991). Because of the apparent high dietary overlap and morphological similarity among syntopic *Carollia* species, they are ideal for testing adaptive and non-adaptive hypotheses, including how bat traits may structure *Piper* consumption and, its converse, how *Piper* traits structure frugivore consumption. At La Selva in Costa Rica, three species of syntopic *Carollia* feed on at least a dozen *Piper* species, with *C. perspicillata* being the most generalist frugivore, *C. sowelli* being intermediate, and *C. castanea* being the most specialized on *Piper* (Fleming 1991). While behavioral studies of *Carollia* bats suggest adaptations to *Piper* scent cues (Thies et al. 1998; Leiser-Miller et al. 2020), broad dietary overlaps among the three bat species imply little specialization, leaving little room for differential frugivore adaptation (Maynard et al. 2019).

**Figure 1.**
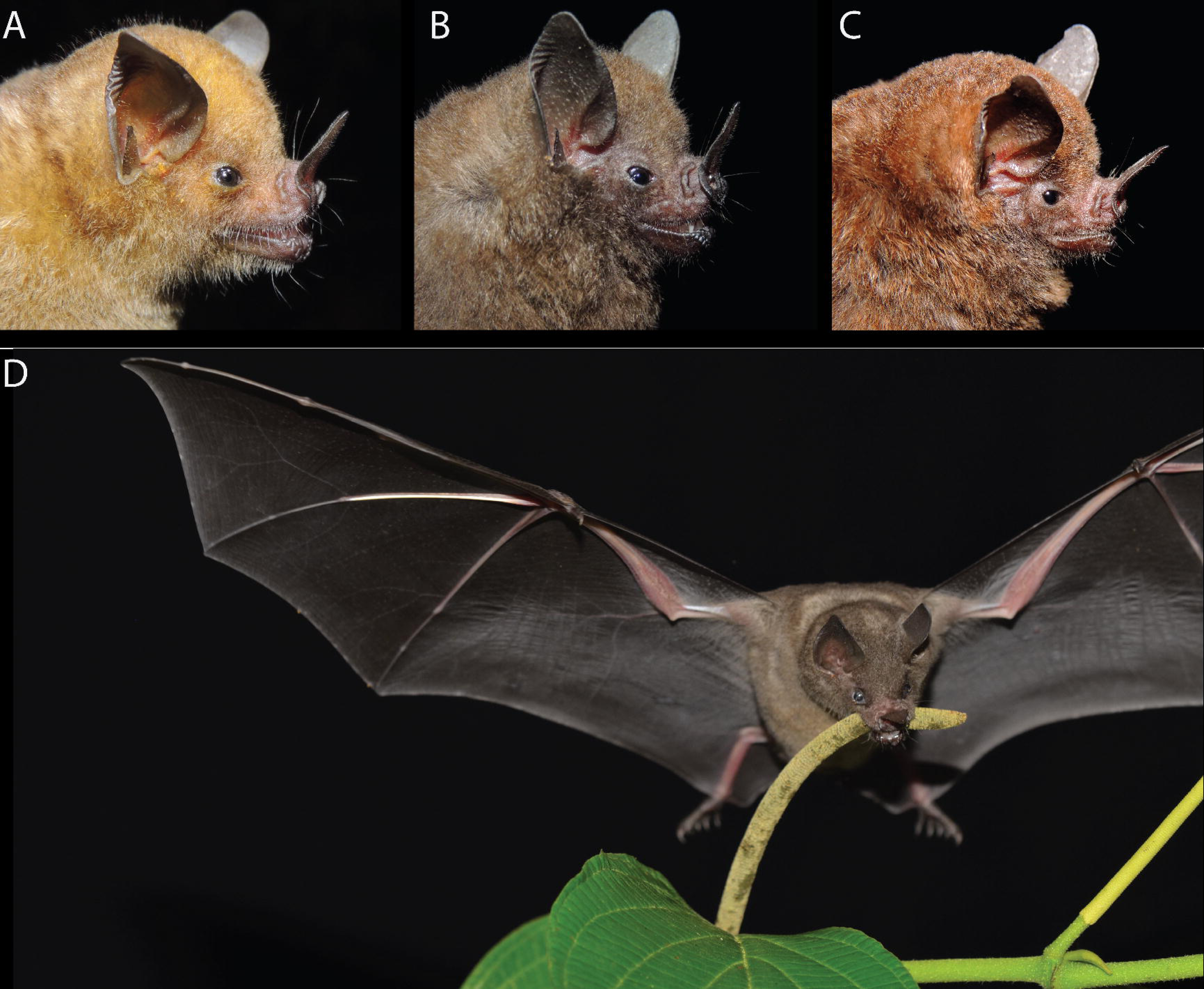
Headshots of the three sympatric short-tailed fruit bats (*Carollia*) found in our study locality in Costa Rica: (A) *Carollia perspicillata*, (B) *C. sowelli*, and (C) *C. castanea*. (D) *C. perspicillata* feeding on *Piper sancti-felicis*. Photo credit: David Villalobos Chaves (A-C) and Susan Whitehead (D).

To evaluate these competing explanations, we conducted a detailed survey of dietary composition in *Carollia* and developed a modeling framework to simultaneously measure the role of traits of both fruits and bats in structuring their ecological interactions. We use Bayesian generalized joint attribute modeling (GJAM) to estimate the consumption indices—an indication of relative consumption rates—of three syntopic species of *Carollia* for *Piper* species, as well as the influence of bat traits on these estimates. In turn, we relate *Piper* fruit traits to these estimates, testing their influence on fruit consumption by bats. Analyzing the trophic interactions among bats and plants, and among competing congeners, requires the integration of several types of ecological data (*e.g*., continuous traits, presence/absence of food resources), and has been historically challenging to model (Clark 2016; Clark et al. 2017). Joint attribute modeling is able to account for multiple sources of variation and multiple predictors of different data types to obtain robust estimates of responses (Clark et al. 2017). In support of the adaptive hypothesis, we predicted that *Piper* traits would reflect dispersal syndromes and therefore relate to fruit consumption indices, and that differential fruit consumption indices would be associated with bat traits. We discovered bat species identity and functional traits structure the consumption of different *Piper* species in the diet, consistent with specialization and niche partitioning, but the *Piper* traits examined showed no relationship to bat-*Piper* consumption indices, indicating those are unlikely to be involved in fruit selection by bats. Despite only subtle trait differences among the bat species studied, our analyses uncovered key differences in consumption contributing to frugivore niche partitioning and therefore adaptation.

## MATERIALS AND METHODS

To test whether traits of *Carollia* bats and *Piper* plants reflect a potentially adaptive match to patterns of interactions between, we collected data from co-occurring individuals of bats and plants at La Selva Biological Station, Sarapiquí, Costa Rica. We used these data to buildlJthree types of Bayesian models. The first models link bats and their traits to *Piper* species represented in bat feces, generating a set of coefficients that describe how each bat trait or species designation shapes the relative consumption tendency for each *Piper* species. We call these modeled coefficients of bat species and traits “*Piper* consumption indices”. The second model estimates the relationship between bat morphometric (*e.g.*, body size) and performance (*i.e.*, bite force) traits related to feeding, and the third quantifies the effects of *Piper* traits on modeled *Piper* consumption indices by each bat species.

### Piper *Consumption by Bats*

To determine how *Carollia* species and traits relate to the consumption of different *Piper* species, we quantified the diets of the three syntopic *Carollia* species at La Selva. All procedures for bat capture and handling were approved by the Institutional Animal Care and Use Committee (IACUC) of the University of Washington, Seattle, USA (protocol #4307-02). We used mist nets to capture bats between 1800-2200 h along trails throughout the forest during the wet season, when there is a greater incidence of fruiting peaks for *Piper* (July and September - December 2015). We collected fecal samples from 318 individuals from the three *Carollia* species (Fig. 1): *C. perspicillata* (*N* = 84), *C. sowelli* (*N* = 111), and *C. castanea* (*N* = 123) by placing individual bats in cloth bags for up to two hours. If the bat defecated, we collected fecal pellets, which we dried in an air-conditioned room for 1-2 days. Samples were then transported to UW for seed identification. We identified seed species in rehydrated fecal pellets using morphological characters and by comparison to a seed reference library that included *Piper* and non-*Piper* species native to La Selva. The reference library was built from seeds removed from ripe fruits collected directly from the parent plant, and plants were identified by LBM, ZAK, Orlando Vargas (OTS), and confirmed via genetic markers (see (Santana et al. 2021)). If we could not identify the species of a particular seed, we classified them as a morphotype (*e.g.*, *Piper* Type 1). We coded each plant species as present or absent in the individual fecal sample (Data S1).

### Bat Traits

We recorded age class (adult, sub-adult, juvenile), sex (male, female), reproductive condition (reproductive, non-reproductive), mass, and forearm length (Data S1) for each bat that produced a fecal sample. Using these bat-specific variables as covariates, we built a model to estimate *Piper* consumption indices, which describe the relationship between bat traits or bat species designation and the probability that a given *Piper* species will be represented in the feces (i.e., to examine how bat traits and species designation influence their dietary records). Our data set was composed of multiple data types, including a zero-inflated matrix of *Piper* species in the bat fecal samples (e.g., 0 if *Piper* species is not present; 1 if *Piper* species is present) and correlates of those data: discrete categories of *Carollia* species, continuous bat size traits, as well as the categorical traits of sex and reproductive condition. Simultaneously estimating relationships among bat species, their traits, and the *Piper* species consumed by bats, is a challenge to general linear models. We implemented the flexible framework of generalized joint attribute modeling (GJAM) (Clark et al. 2017), which uses a Bayesian multivariate approach to infer the parameters of the linear model based on a series of joint distributions of both the bat traits and the *Piper* fecal abundances, while simultaneously accommodating multifarious trait data, in this case from bats.

### Generalized Joint Attribute Modeling

For each observation i of n bat individuals, there is a set {x_i_, y_i_ }^n^, in which each x_i_ observation has Q predictors to result in a vector of predictors x_iq_:1 … Q. In our case Q = 6, with predictors species, age class, sex, reproductive condition, mass, and forearm length. The set of responses is a vector of y_ip_:1 … P, where P is the total number of *Piper* species (P = 18) observed across all fecal samples. For y_ip_, each vector of bat individual i is the presence or absence of *Piper* species p. Seven *Piper* species were removed from the analysis, as they accounted for less than 1% of the observations (Fig. S2). Most of the observations in y_ip_ are 0, meaning most *Piper* species are not observed in a sample. To accommodate this zero-inflation, GJAM implements a Tobit regression. The representations of x_i_ and y_i_ are composed of partitions of discrete and continuous space, and GJAM applies a connection between the two, which we represent as I in our model. Thus, it is possible to estimate a continuous response w_i_ from multifarious data such that for each observation,

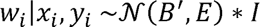

where B’ is the matrix of coefficients and E is a P x P correlation matrix to represent the covariances among the response variables. For detailed explanations of the calculations of I, E, and w, see further discussion in (Clark et al. 2017). We estimated the coefficients using the R package gjam v. 2.1.6 for 20,000 generations, discarding 4,000 as burn-in. We applied a series of dimension reduction options (*N* = 2, 5, by *r* = 2, 5) to facilitate convergence amidst the multiple dimensions of covariance space and adopted the one that yielded the lowest model deviance. Note that we compared both fractional composition models (continuous on (0,1)) and presence-absence models (discrete). Medians of the posterior distributions of the continuous response w_i_ were used for further modeling.

### Bat Functional Traits

Bite force is a metric of feeding performance linked to the mechanical demands of the food a species can process (Aguirre et al. 2002; Santana et al. 2010; Santana and Miller 2016). Following methods by (Santana et al. 2010), we measured deep bilateral, voluntary bite forces for at least ten wild individuals per *Carollia* species using a piezoelectric force transducer (Kistler 9203; range ±500 N, accuracy 0·01–0·1 N) attached to a handheld charge amplifier (Kistler 5995A). The force transducer was mounted between two metal plates covered with medical tape to provide a non-skid biting surface and to protect the bats’ teeth. We adjusted the distance between the bite plates for each individual to accommodate a moderate gape angle of approximately 30°, following (Santana et al. 2010). To avoid variation from age (Santana and Miller 2016) and stress to reproductive females, we only measured adult males and adult non-pregnant, non–lactating females. We recorded five to eight measurements for each bat and chose the highest value to represent maximum bite force. Following bite force measurements, we recorded head length, width, and height measured to the nearest 0.1 mm (Fig. S1B), as well as mass and forearm length for most individuals (Data S1).

### Piper *Fruit and Seed Traits*

Physical traits of fruits and seeds can constrain whether and how bats of different sizes can process them. We collected dimensions of whole *Piper* infructescences (the unit consumed by *Carollia*, called “fruits” throughout this paper for simplicity) and individual seeds to estimate how these traits relate to the modeled *Piper* consumption indices. We measured length and width from five ripe fruits from each *Piper* species to the nearest 0.001 mm, and used ImageJ (Rasband, W.S., ImageJ, U. S. National Institutes of Health, Maryland, USA) to measure seed length and seed width from digital photographs of three seeds from each fruit. Seed photos were taken with a Leica MZ 95 microscope camera coupled with Clemex Captiva software. We used these fruit and seed measurements to calculate a ratio (length/width) as an estimate of fruit and seed shape, respectively.

### Bayesian Hierarchical Modeling

After determining that both sex and head length were linear predictors of bite force in regressions with either a sample-wide intercept (male sex coefficient *t*_(27)_ = 2.29, *P* -value = 0.03, head length coefficient *t*_(27)_ = 7.60, *P* -value = 3.54e-08), or species-specific intercepts (male sex coefficient *t*_(27)_ = 4.23, *P* -value 1.20e-04, head length coefficient *t*_(27)_ = 2.44, *P* -value = 0.01), we modeled bite force as a function of bat body size traits while controlling for both sex and head length, which may explain bite force. We used Jags v.3.3.0 (Plummer 2003) to code these models, and ran them in the R package R2jags v.0.04-01 (Su and Yajima 2012). These models included species-specific intercepts whose prior was drawn from a normal distribution. Priors for both between- and within-population variances were modeled as half-Cauchy distributions with variance of at least 100,000. These priors do not make any assumptions about the relative contribution of variation from different levels in the hierarchy (Gelman and Hill 2006). For each model, four independent chains ran for 500,000 iterations with 250,000 iterations as burn-in, and samples were taken every 250 generations. Convergence was assessed by both the effective sampling size of model parameters (>1000 in every case), and the potential scale reduction factor (PSRF), which approaches 1 at convergence (Gelman and Rubin 1992). The models coded measures of error to estimate the variance explained, as outlined by (Gelman and Pardoe 2006).

We used the *Piper* traits as regressors in Bayesian models of *Piper* consumption indices by bat estimated by GJAM analysis. Thus, these models connect the differential use of *Piper* resources by bats (e.g., across species, age class, or body sizes) to the *Piper* traits that might underlie those differences. We used the R package MCMCglmm (Hadfield 2010) to code the models, and accounted for the correlation structure of the data due to evolutionary relatedness by including a molecular phylogeny of *Piper* (Santana et al. 2021) as a species-specific (random) effect. We applied a parameter-expanded prior with the parameters V = 1 nu = 1 for the residual variance (Rojas et al. 2018), and a proper Cauchy prior defined by V = 0.5 nu = 1 and alpha.mu = 0 and alpha.V = 10^3^ for the random term (Hadfield 2019). Each model ran for 200,000 iterations, sampling every 100, with 10,000 generations as burn-in. Convergence of the resulting posteriors was assessed by the effective sampling size of model parameters (>1000 in every case). In total, we ran four models corresponding to the modeled bat *Piper* consumption indices associated with bat forearm, body mass, and *C. castanea* and *C. perspicillata*.

## RESULTS

As expected, the percentage of *Piper* presence in the diet was highest in the specialist *C. castanea* (67.5%) and lowest in the generalist *C. perspicillata* (45.2%). *C. sowelli* was intermediate (60.5%) (Fig. S2). For the period sampled, *Piper* Type 4 was the most common species in the diets of *C. castanea* and *C. sowelli*, while *P. hispidum* was the most common for *C. perspicillata*. Dietary proportions are displayed in Fig. S2 and raw diet data in Data S1.

### Piper *consumption indices across different* Carollia

Model fit was assessed through DIC (Deviance Information Criterion) and posterior predictive output and the fractional composition model (as opposed to presence-absence) demonstrated a much better fit (Fig. 2A, 2B). In GJAM, the sensitivity of the model to various covariate inputs (*i.e.*, the bat traits and species designations) can be interpreted as the amount of information each input contributes to estimating the model coefficients (Clark et al. 2017). A positive consumption index in a bat species indicates greater relative consumption of that Piper species given a particular covariate, while a negative consumption index indicates the opposite. Bat species (particularly *C. castanea* and *C. perspicillata*), forearm length, age, and reproductive status all showed sensitivity values greater than one, suggesting they were much more informative than sex or body mass in explaining the presence of *Piper* species in the diet of *Carollia* (i.e., *Piper* bat consumption indices; Table S1; Fig. 2C). Figure 3 illustrates the Piper consumption indices for each bat species, *i.e.*, the posterior probabilities for each *Piper* species, estimated by the consumption index of each bat species for that particular *Piper* species (details in Table S2). While the 95% highest posterior density (HPD) credible interval crossing zero corresponds to a weak relationship between the covariate and the *Piper* species, an HPD not overlapping zero can be interpreted as a strong response. Consequently, six species of *Piper* showed a strong positive response to *C. perspicillata* (in order of highest consumption index: *P. hispidum* (median: 0.45; 95% HPD: [0.22, 0.66]), *P*. *colonense* (0.45 [0.20, 0.66]), *P. silvivagum* (0.45 [0.20, 0.68]), Type 4 (0.36 [0.13, 0.56]), *P. aduncum* (0.35 [0.10, 0.59]), and Type 10 (0.31 [0.001, 0.59])). The *Piper* specialist *C. castanea* also has the lowest consumption indices for five of these six species (Fig. 3); *P. colonense* (-0.52 [-0.75, -0.24])*, P. hispidum* (-0.34 [-0.56, - 0.09]), *P. silvivagum* (-0.34 [-0.58, -0.07]), *P. aduncum* (-0.33 [-0.58, -0.07]), and Type 4 (-0.32 [-0.54, -0.08]). *C. castanea* also showed a negative consumption index for *P. sancti-felicis* (-0.33 [-0.63, -0.01]), towards which *C. sowelli* (the bat species that exhibits intermediate specialization on *Piper*) also demonstrated a positive consumption index (0.22 [0.02, 0.41])*. C. perspicillata* only showed a negative consumption index towards *Piper* Type 1 (-0.29 [-0.58, -0.01]). Table S2 shows coefficient estimates for all *Piper* species.

**Figure 2.**
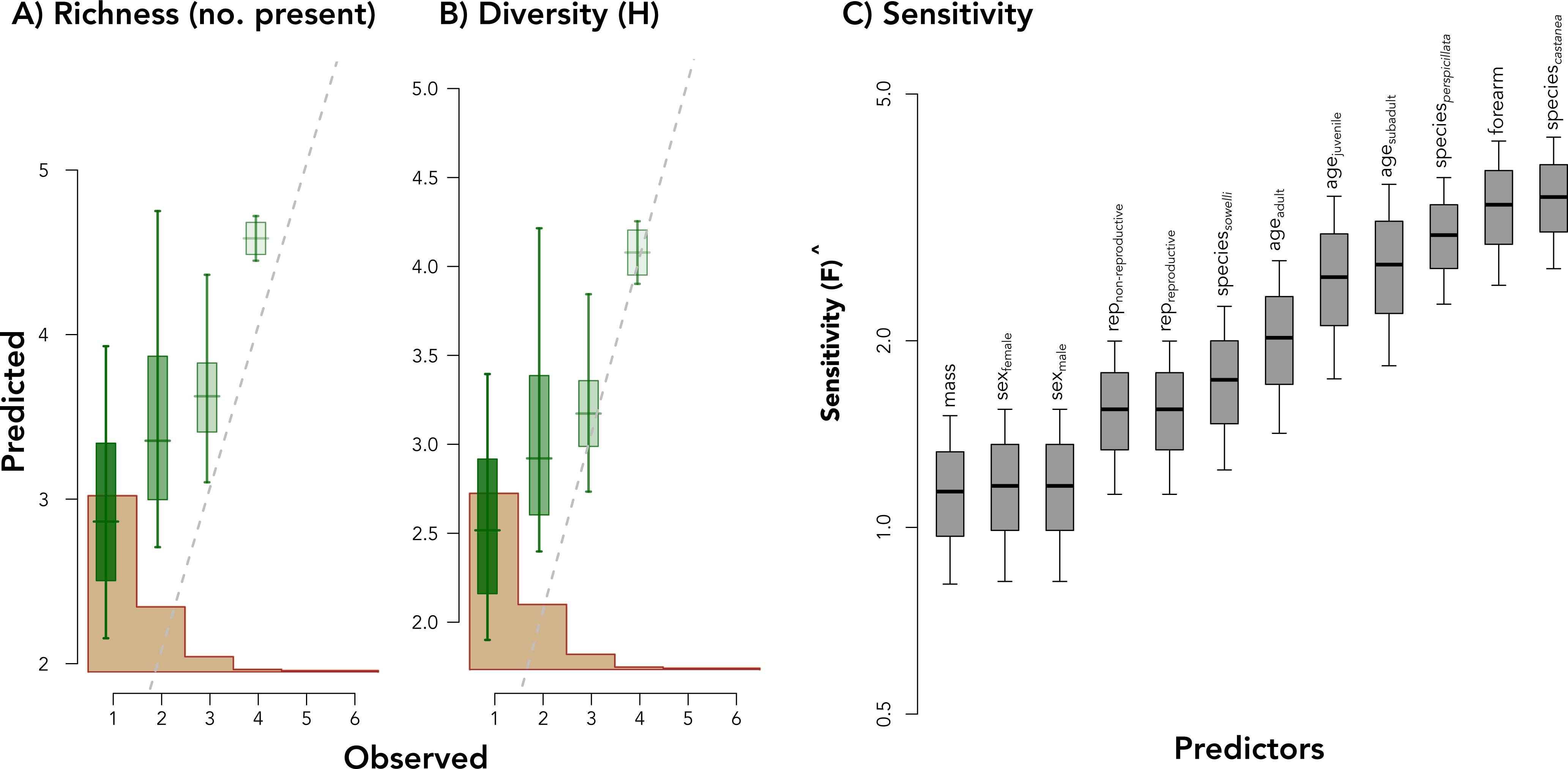
Parameters (A) richness and (B) diversity of *Piper* species per fecal sample calculated as posterior predictive checks of model fit for the GJAM relative abundance fractional composition model. Model fit of the generalized joint attribute modeling in predicting *Piper* consumption indices. The brown histogram in (A) and (B) is the distribution of the observed data and the dashed lines are the 1:1 diagonals of the observed values and predictions. (A) Richness represents a predictive posterior check, such that richness is responses predicted that are greater than 0. (B) More diverse “sites” had a better fit, as less diverse “sites” are more rare. (C) Sensitivity of the model to covariate inputs can be interpreted as the amount of information each input contributes overall to estimating the model coefficients. The higher the sensitivity, the more informative the covariate to the model.

**Figure 3.**
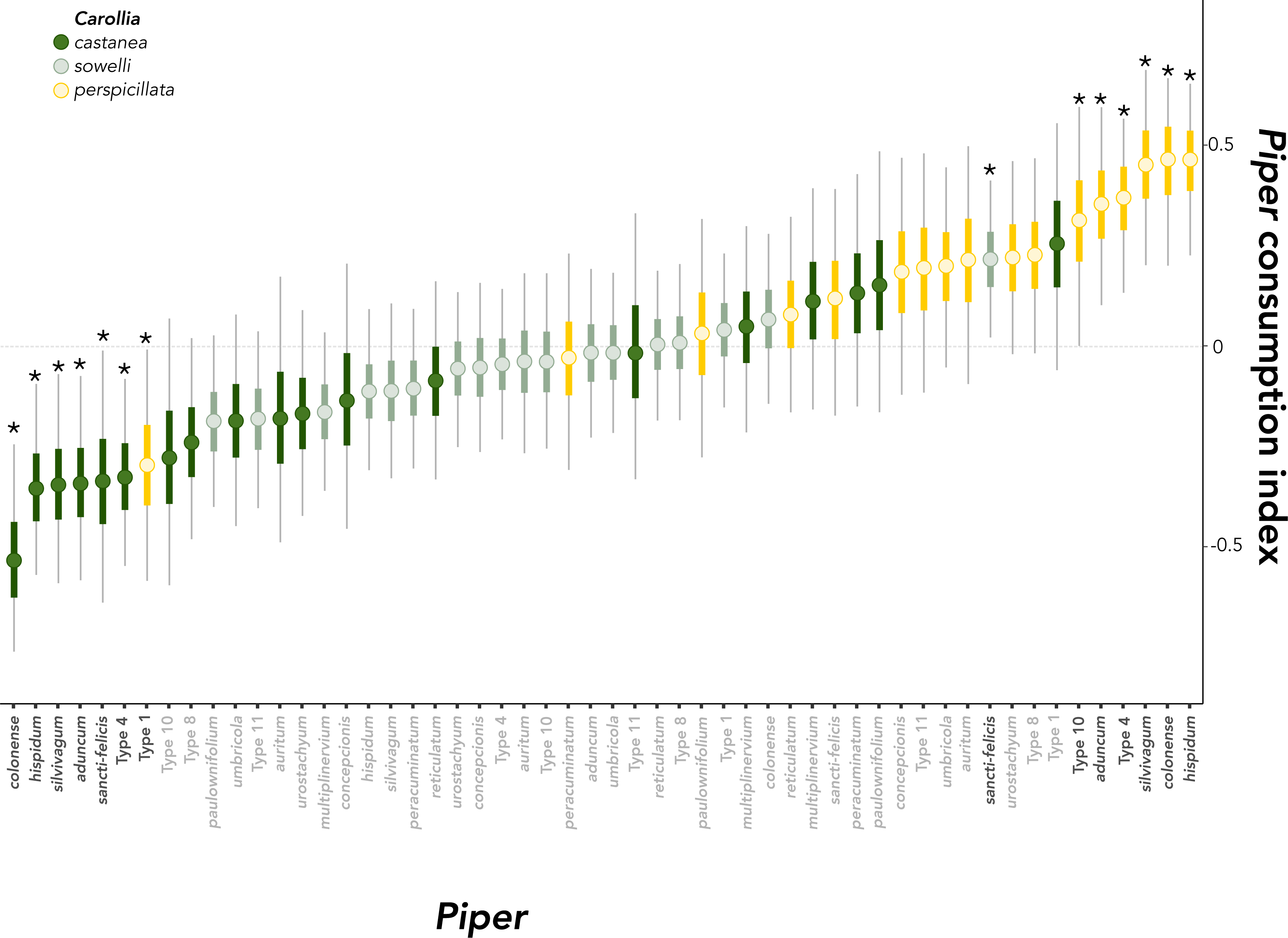
Posterior distributions of model coefficients (*Piper* consumption indices per bat species), ordered by median. *Piper* consumption indices can be interpreted as the probability of a particular *Carollia* species to show a higher or lower consumption index for a particular *Piper* species. Asterisks and black species names refer to *Piper* in which 95% of the highest posterior density intervals did not cross zero, indicating a strong positive or negative response.

### *Influence of bat traits on* Piper *consumption indices*

The sensitivity of the model to the bat traits used as model inputs and their influence on consumption indices of *Piper* species varied (Table S1; Fig. 2C). The magnitude of these coefficients reflects the influence of the trait on the consumption index, or overall patterns of *Piper* consumption. Sensitivity was high for one covariate representing body size (forearm length), which strongly influenced the consumption indices for several *Piper* species (Fig. 2C; Fig. 4). The consumption index distribution for *P.* Type 1 showed the strongest positive response to forearm length (0.70 [0.34,1.01]). There was a strong positive influence of forearm length in four other *Piper* species (Fig. 4; *P. peracuminatum*: 0.41 [0.13, 0.73]; *P. paulowniifolium*: 0.34 [0.04, 0.70], *P. sancti-felicis*: 0.27 [0.01, 0.59], and *P. multiplinervium*: 0.26 [0.03, 0.52]. A strong negative response to forearm was estimated for *Piper* Type 4 (-0.21 [- 0.37, -0.06]), which also had anticorrelated consumption indices favoring *perspicillata* (0.36 [0.13, 0.56]) and negative for *castanea* (-0.31 [-0.54, -0.08]). Although age showed the second highest sensitivity among all covariates (Fig. 2C), no *Piper* species had a posterior that entirely excluded zero (Fig. S4), likely because there were few observations of juveniles and subadults. It is worth noting that despite this variation, *P. paulowniifolium* showed the strongest response with adult bats and Type 1 and *P. hispidum* showed the strongest response in juveniles and subadults (Fig. S4). There was no meaningful bat sex or reproductive condition influence on *Piper* species consumed. Table S2 summarizes estimates for each categorical or continuous covariate of this model.

**Figure 4.**
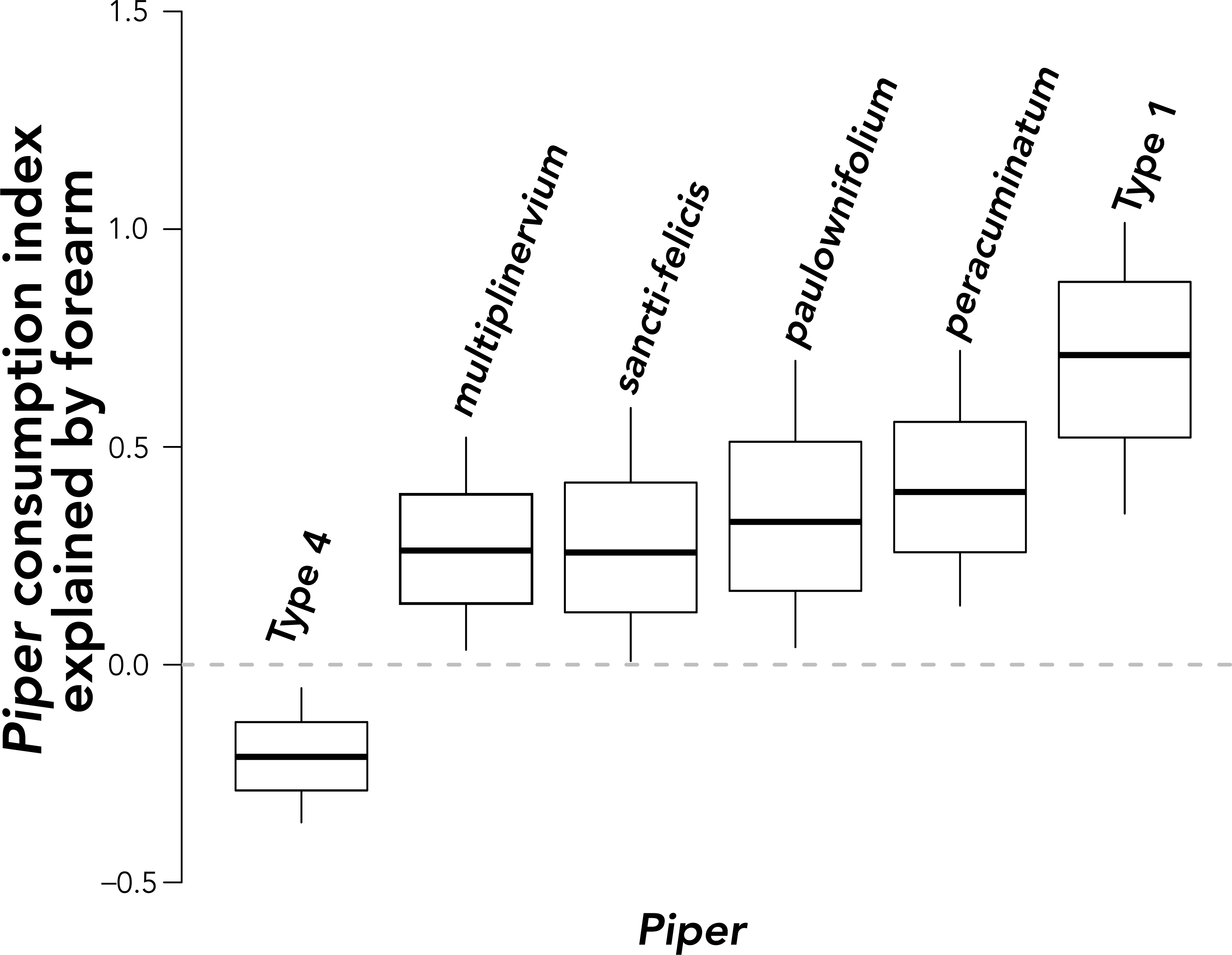
Posterior distributions of model coefficients (Piper consumption indicies) in response to forearm length. Coefficients are only shown for *Piper* species that indicate strong positive or negative responses, determined by the entire 95% highest posterior density being entirely above or below zero.

### Bat functional traits

We modeled the scaling of bite force with head and body dimensions using hierarchical models and in the natural log scale in every case. Although head length did not differ among species (*F* _2,27_ = 1.443, *p* = 0.256), it was a positive covariate of maximum bite force with high variance explained (multiple regression *R* ^2^ = 0.90, after controlling for sex), and a consistently positive posterior coefficient distribution (Table S3). Similar results were obtained in combination with body mass (multiple regression *R* ^2^ = 0.90), and forearm length (multiple regression *R* ^2^ = 0.90 and lowest deviance), with the forearm length coefficient indicating negative trends with bite force after controlling for head length (Table S4). Head and forearm length were positively correlated (*R* = 0.76, *t*_28_ = 6.2306, *p* = 9.862e-07). In short, once the effect of head size is accounted for, and acknowledging that larger bats have larger heads, the marginal relationship between forearm length and bite force tends to be negative. Male bats always had greater bite force compared to females, even after controlling for head length, or body size (Table S3).

### Piper *fruit traits and Bat*-Piper responses

Phylogenetic hierarchical Bayesian models sought to relate *Piper* consumption indices per bat species to *Piper* traits (seed shape index, fruit shape index). These models examined whether *Piper* traits could predict the relative strength of GJAM coefficients reflecting the likelihood that a given bat will consume a given *Piper* species (*i.e.*, *Piper* bat consumption indices). Neither of the fruit traits was a statistically significant predictor of *C. castanea* consumption indices (Table S4), or of consumption indices estimated based on forearm length or body mass for *C. perspicillata*. *C. sowelli* did not have any outlier consumption indices.

## DISCUSSION

Through adaptation, plant-animal interactions may result in the reciprocal influence of fruits on frugivory and vice versa, suites of matching traits on both sides of the mutualism. However, it is only when ecological traits are measured in both fruits and frugivores simultaneously while considering within-species variation that one can infer whether measured traits in plant-animal mutualisms are congruent with this scenario. We overcame these challenges by using generalized joint attribute modeling that modeled the occurrence of *Piper* in bat diets while considering multiple covariates with different variance structures simultaneously. By modeling the influence of both bat and fruit traits on the interaction, we tested whether plant and bat traits predict the structure of bat dietary composition. Our finding that differential *Piper* fruit consumption is driven by bat species and their traits, primarily body size, yet no support for fruit traits influencing bat consumption supports an asymmetric influence of fruit on frugivore specialization but not its converse.

Consumption of several *Piper* species is non-random, and strongly predicted by the identity of the bat species and forearm length, a strong covariate of body size in bats. Species identity primarily influenced consumption indices, with the syntopic *Carollia castanea* and *perspicillata* at opposite ends of consumption index variation. Of the six *Piper* species with the highest consumption index by the generalist *C. perspicillata*, five also had the lowest consumption index by the specialist *C. castanea* (Fig. 3). In contrast, previous work found species identity influenced the proportion of *Piper* in fecal samples, but did not affect *Piper* dietary composition by individual bat and as a result there was near-complete dietary niche overlap among the three bat species (Maynard et al. 2019). In those prior analyses, the relationship between species identity and traits was estimated by relating each variable of interest (e.g., traits such as species identity, sex, age) to distances obtained through non-metric multidimensional scaling ordination of *Piper* abundance using generalized additive models.

Finding an inverse consumption index for a suite of *Piper* species is evidence that *Carollia* bats do partition *Piper* resources, contrary to previous results. Our results suggest GJAM models achieve greater sensitivity by allowing for simultaneous inference of multiple covariates with different variance structures, helping elucidate the patterns of species interaction within this guild, capturing the richness of the samples (Fig. 2). While *C. perspicillata* has a more flexible diet that includes many non-*Piper* fruits (Fig. S2) or even nectar and insects, when it eats *Piper*, it uses *Piper* species that *C. castanea* uses seldomly. This indicates these bats partition the dietary niche in previously unsuspected ways. In line with previous results, however, nearly all other *Piper* species had overlapping consumption indices for the three bat species (Fig. 3), indicating dietary niche overlap among *Carollia* bats for most but not all *Piper* species.

Besides species identity, body size as measured by forearm length also structured consumption indices for some *Piper* species. Consumption indices for *Piper* Type 1 and Type 4 separate *C. castanea* and *C. perspicillata*, and for *P*. *sancti-felicis* separate *C. castanea* and *C. sowelli*. These *Piper* species also showed a strong response to bat forearm length, even after accounting for bat species identity. Differences in body size that structure *Piper* consumption indices may indicate differences in dietary niche breadth because niche breadth can increase with body size in bats via larger home ranges (Barclay and Brigham 1991). Instead of specialization, *Piper* consumption indices might be related to *Piper* geographic distribution and bat dispersal ability. In effect, and although we did not focus on non-*Piper* species, the larger generalist *C. perspicillata* eats fruits from several other types of plants too. Relating niche breadth to body size would thus support ongoing competition among bat congeners. These two bat species may also use their habitat differently or at different times or be in active competition on an ecological time scale. We propose that in the presence of a competing species such as *C. perspicillata*, the realized niche is smaller for the specialist *C. castanea,* such that it specializes on different *Piper* resources reduces niche overlap. In terms of relating to the functional ecology of body size differences, structuring of *Piper* consumption indices by size —which predicts bite force (Table S3)— aligns our results with comparative analyses for all phyllostomids in which bite force relates to consumption or larger and/or tougher fruit (Santana et al. 2010). While our link between bite force and *Piper* consumption is indirect, our results further support an adaptive hypothesis for bat traits on fruit consumption.

Under an adaptive scenario and dispersal syndrome hypothesis, a reciprocal association between frugivore phenotype and traits of the food resources is expected in the context of coevolution in plant-animal mutualisms (Valenta and Nevo 2020), as in the case of beak size and shape, and seed size and hardness in Galapagos finches (Schluter and Grant 1984; Schluter et al. 1985). We found no relationship between *Piper* traits and any consumption indices estimates, suggesting *Piper* fruit morphologies are not adaptive to signaling specific *Carollia* frugivores. While there is empirical evidence of fruit morphologies correlating with traits of their dispersers (Janson 1983; Valenta and Nevo 2020), our results suggest morphological traits of the animal disperser likely structure this particular mutualism. In an asymmetry that detracts from the dispersal syndrome co-evolutionary hypothesis, morphological traits of the frugivore appear to be shaped by *Piper* consumption but not the other way around. For plants, slower evolutionary rates in outbreeding populations (Herrera 1984; Valenta and Nevo 2020), and generalism may explain this asymmetry. As with many animal-dispersed fruiting plants, *Piper* is also consumed by other non-bat frugivores (e.g., birds) (Palmeirim et al. 1989) for which seed and fruit morphology may play a role.

Unmeasured fruit traits might also be selectively shaped in this plant-bat interaction. Traits such as fruiting time (Thies and Kalko 2004), plant habit, or secondary metabolite profiles (Whitehead et al. 2016; Santana et al. 2021) have been proposed as being more important to differential consumption than the physical traits of fruit we measured. There is also strong support for chemical communication between plants and bats, with behavioral evidence for *Carollia* using the sense of smell to locate ripe fruit (Thies et al. 1998; Leiser-Miller et al. 2020), and bat olfactory receptor diversity scaling to dietary diversity (Yohe et al. 2021). Chemical bouquet composition both differs sharply and evolved adaptively among *Piper* species (Santana et al. 2021), so those traits may affect and better reflect reciprocal adaptation to bat consumption. In short, while we found no evidence of an effect of fruit and seed dimensions on bat consumption, behavioral and chemical evidence suggest scent traits are likely to be more important in structuring niche partitioning across bat species.

Though the inverse relationship of consumption indices for *Piper* may indicate ongoing specialization in food resources in two *Carollia* species, there is some indication that behavioral aspects, such as learning, also contribute to differential resource use. While no *Piper* species showed a significant association to bat age, age had high sensitivity and some interesting patterns of contrasting consumption tendencies in adult versus juvenile bats warrant further exploration (Fig. S3). A previous study has found that adults used a lower percentage of mid-to late-successional species than juveniles, partitioning *Piper* by habitat (Maynard et al. 2019). We hypothesize that older, more experienced bats can locate and exploit resources better than younger, naïve bats –whether through spatial learning, or familiarity with less conspicuous fruit cues.

Because our model both accounts for several sources of variation and can incorporate many different types of ecological data, we were able to discover partitioning and estimate the influence of various traits on plant-frugivore interactions. Identifying such patterns provided quantitative evidence of the relationships between differential resource use and frugivore traits. We discovered that, while the use of different fruit resources is related to putatively adaptive differences in body size traits, age may also play an important role in defining the dietary niche of syntopic species. As body size both may confer niche breadth and underlies functional traits such as bite force, our findings are consistent with both specialization through adaptation and ongoing competition among bat frugivores. While there was not an effect of the plant traits examined on bat consumption, mounting evidence for plant chemical adaptation and specialization in this system suggests plant-bat interactions may not be mediated by gross fruit morphology. Thus, this approach enabled us to both uncover the most informative predictors of differential plant use, and hint at new mechanisms underlying the evolutionary ecology of fruit-frugivore interactions.

## Acknowledgements

This project was made possible by the scientists and administrators at La Selva Biological Research Station in Sarapiqui, Costa Rica. We specifically acknowledge Bernal Rodriguez Herrera, Bernal Matarrita, Orlando Vargas, David Villalobos Chaves, and Minor Porras for their technical assistance and help with permits. This project was funded via the National Science Foundation DEB-1442142, DEB-1456375, DEB-1856776, and the Postdoctoral Research Fellowship in Biology (NSF DBI-1812035).

