## Supplementary Information for "Testing for reciprocal trait influence in plant-frugivore interactions using generalized joint attribute modeling"

**Table S1**. Sensitivity estimates for each covariate response to *Piper* plant in GJAM model, corresponding to Figure 2C.

| Variable | Sensitivit y | Standard Error |
|---|---|---|
| $Bat_{castanea}$ | 2.93 | 0.384 |
| $Bat_{sowelli}$ | 1.65 | 0.245 |
| $Bat_{perspicillata}$ | 2.75 | 0.345 |
| Forearm | 3.15 | 0.443 |
| Mass | 1.32 | 0.191 |
| $Sex_{female}$ | 1.06 | 0.166 |
| $Sex_{male}$ | 1.06 | 0.166 |
| $Age_{adult}$ | 1.91 | 0.305 |
| $Age_{subadult}$ | 2.60 | 0.433 |
| $Age_{juvenile}$ | 2.45 | 0.407 |
| $Rep_{non-repreproductive}$ | 1.42 | 0.197 |
| $Rep_{reproductive}$ | 1.42 | 0.197 |

**Table S2**. Coefficients of response variables for each *Piper* plant species. Estimates are mean prediction values for each variable. SD is posterior standard deviation. The percentage columns are the upper and lower boundary of the 95% credible interval of the posterior distribution. Rows highlighted in **bold** are estimates in which the entire 95% credible interval does not cross zero, the +/− indicating the direction of this estimate.

| *Piper* | variable | Estimate | StdErr | 2.5% | 97.5% |
|---|---|---|---|---|---|
| *aduncum* | **$\text{Bat}_{\textit{castanea}}$** | **−0.333** | **0.128** | **−0.575** | **−0.072** |
| | **$\text{Bat}_{\textit{perspicillata}}$** | **0.348** | **0.124** | **0.102** | **0.589** |
| | $\text{Bat}_{\textit{sowelli}}$ | −0.015 | 0.106 | −0.224 | 0.191 |
| | $\text{Sex}_{\text{male}}$ | 0.062 | 0.074 | −0.083 | 0.207 |
| | $\text{Age}_{\text{juvenile}}$ | −0.165 | 0.197 | −0.553 | 0.215 |
| | $\text{Age}_{\text{subadult}}$ | 0.143 | 0.202 | −0.250 | 0.542 |
| | $\text{Rep}_{\text{reproductive}}$ | 0.022 | 0.142 | −0.255 | 0.299 |
| | mass | −0.045 | 0.088 | −0.218 | 0.129 |
| | forearm | 0.055 | 0.114 | −0.155 | 0.294 |
| *auritum* | $\text{Bat}_{\textit{castanea}}$ | −0.172 | 0.167 | −0.482 | 0.172 |
| | $\text{Bat}_{\textit{perspicillata}}$ | 0.210 | 0.150 | −0.092 | 0.493 |
| | $\text{Bat}_{\textit{sowelli}}$ | −0.038 | 0.113 | −0.262 | 0.180 |
| | $\text{Sex}_{\text{male}}$ | 0.016 | 0.078 | −0.137 | 0.168 |
| | $\text{Age}_{\text{juvenile}}$ | −0.066 | 0.198 | −0.454 | 0.322 |
| | $\text{Age}_{\text{subadult}}$ | 0.000 | 0.210 | −0.412 | 0.419 |
| | $\text{Rep}_{\text{reproductive}}$ | 0.066 | 0.143 | −0.212 | 0.348 |
| | mass | −0.083 | 0.091 | −0.262 | 0.091 |
| | forearm | 0.249 | 0.172 | −0.059 | 0.606 |
| *colonense* | **$\text{Bat}_{\textit{castanea}}$** | **−0.520** | **0.133** | **−0.750** | **−0.241** |
| | **$\text{Bat}_{\textit{perspicillata}}$** | **0.452** | **0.120** | **0.199** | **0.660** |
| | $\text{Bat}_{\textit{sowelli}}$ | 0.067 | 0.107 | −0.141 | 0.277 |
| | $\text{Sex}_{\text{male}}$ | −0.080 | 0.073 | −0.225 | 0.063 |
| | $\text{Age}_{\text{juvenile}}$ | −0.098 | 0.201 | −0.499 | 0.293 |
| | $\text{Age}_{\text{subadult}}$ | 0.015 | 0.215 | −0.407 | 0.436 |
| | $\text{Rep}_{\text{reproductive}}$ | 0.082 | 0.146 | −0.204 | 0.367 |
| | mass | 0.008 | 0.086 | −0.160 | 0.177 |
| | forearm | −0.052 | 0.103 | −0.240 | 0.162 |
| *concepcionis* | $\text{Bat}_{\textit{castanea}}$ | −0.129 | 0.167 | −0.448 | 0.204 |
| | $\text{Bat}_{\textit{perspicillata}}$ | 0.181 | 0.149 | −0.119 | 0.464 |
| | $\text{Bat}_{\textit{sowelli}}$ | −0.052 | 0.107 | −0.259 | 0.157 |

| | | | | | |
|---|---|---|---|---|---|
| | $Sex_{male}$ | 0.065 | 0.078 | −0.086 | 0.218 |
| | $Age_{juvenile}$ | −0.096 | 0.200 | −0.492 | 0.292 |
| | $Age_{subadult}$ | −0.022 | 0.214 | −0.442 | 0.392 |
| | $Rep_{reproductive}$ | 0.117 | 0.146 | −0.169 | 0.408 |
| | mass | −0.160 | 0.088 | −0.334 | 0.011 |
| | forearm | 0.225 | 0.171 | −0.073 | 0.597 |
| *hispidum* | **$Bat_{castanea}$** | **−0.344** | **0.121** | **−0.562** | **−0.092** |
| | **$Bat_{perspicillata}$** | **0.453** | **0.109** | **0.224** | **0.646** |
| | $Bat_{sowelli}$ | −0.110 | 0.101 | −0.304 | 0.092 |
| | $Sex_{male}$ | 0.077 | 0.068 | −0.057 | 0.210 |
| | $Age_{juvenile}$ | −0.151 | 0.174 | −0.497 | 0.188 |
| | $Age_{subadult}$ | 0.278 | 0.185 | −0.092 | 0.634 |
| | $Rep_{reproductive}$ | −0.127 | 0.125 | −0.369 | 0.118 |
| | mass | −0.058 | 0.080 | −0.213 | 0.098 |
| | forearm | 0.097 | 0.097 | −0.085 | 0.298 |
| *multiplinervium* | $Bat_{castanea}$ | 0.114 | 0.140 | −0.155 | 0.389 |
| | $Bat_{perspicillata}$ | 0.047 | 0.130 | −0.211 | 0.296 |
| | $Bat_{sowelli}$ | −0.161 | 0.099 | −0.355 | 0.035 |
| | $Sex_{male}$ | 0.088 | 0.070 | −0.050 | 0.226 |
| | $Age_{juvenile}$ | 0.027 | 0.181 | −0.336 | 0.380 |
| | $Age_{subadult}$ | 0.025 | 0.197 | −0.356 | 0.417 |
| | $Rep_{reproductive}$ | −0.052 | 0.132 | −0.306 | 0.207 |
| | mass | −0.063 | 0.080 | −0.218 | 0.093 |
| | **forearm** | **0.264** | **0.125** | **0.032** | **0.523** |
| *paulownifolium* | $Bat_{castanea}$ | 0.154 | 0.164 | −0.162 | 0.480 |
| | $Bat_{perspicillata}$ | 0.030 | 0.150 | −0.273 | 0.314 |
| | $Bat_{sowelli}$ | −0.184 | 0.108 | −0.395 | 0.028 |
| | $Sex_{male}$ | 0.042 | 0.074 | −0.104 | 0.187 |
| | $Age_{juvenile}$ | −0.069 | 0.187 | −0.440 | 0.296 |
| | $Age_{subadult}$ | −0.096 | 0.205 | −0.499 | 0.306 |
| | $Rep_{reproductive}$ | 0.165 | 0.139 | −0.106 | 0.442 |
| | **mass** | **−0.240** | **0.087** | **−0.413** | **−0.073** |
| | **forearm** | **0.340** | **0.172** | **0.038** | **0.700** |
| *peracuminatum* | $Bat_{castanea}$ | 0.133 | 0.146 | −0.147 | 0.424 |
| | $Bat_{perspicillata}$ | −0.030 | 0.135 | −0.303 | 0.229 |
| | $Bat_{sowelli}$ | −0.103 | 0.100 | −0.300 | 0.093 |
| | $Sex_{male}$ | 0.066 | 0.069 | −0.070 | 0.201 |

| | | | | | |
|---|---|---|---|---|---|
| | $Age_{juvenile}$ | 0.137 | 0.175 | −0.203 | 0.478 |
| | $Age_{subadult}$ | −0.227 | 0.197 | −0.612 | 0.156 |
| | $Rep_{reproductive}$ | 0.089 | 0.128 | −0.160 | 0.341 |
| | **mass** | **−0.213** | **0.079** | **−0.368** | **−0.059** |
| | **forearm** | **0.405** | **0.150** | **0.134** | **0.725** |
| *reticulatum* | $Bat_{castanea}$ | −0.084 | 0.124 | −0.327 | 0.161 |
| | $Bat_{perspicillata}$ | 0.079 | 0.123 | −0.162 | 0.319 |
| | $Bat_{sowelli}$ | 0.005 | 0.093 | −0.182 | 0.186 |
| | $Sex_{male}$ | 0.009 | 0.067 | −0.122 | 0.142 |
| | $Age_{juvenile}$ | 0.106 | 0.177 | −0.249 | 0.445 |
| | $Age_{subadult}$ | −0.035 | 0.195 | −0.413 | 0.350 |
| | $Rep_{reproductive}$ | −0.071 | 0.128 | −0.320 | 0.182 |
| | mass | 0.091 | 0.075 | −0.057 | 0.241 |
| | forearm | −0.010 | 0.088 | −0.180 | 0.166 |
| *sancti-felicis* | **$Bat_{castanea}$** | **−0.329** | **0.157** | **−0.630** | **−0.010** |
| | $Bat_{perspicillata}$ | 0.114 | 0.142 | −0.170 | 0.388 |
| | **$Bat_{sowelli}$** | **0.215** | **0.099** | **0.022** | **0.408** |
| | $Sex_{male}$ | 0.018 | 0.070 | −0.118 | 0.157 |
| | $Age_{juvenile}$ | −0.192 | 0.184 | −0.554 | 0.172 |
| | $Age_{subadult}$ | 0.145 | 0.193 | −0.237 | 0.516 |
| | $Rep_{reproductive}$ | 0.046 | 0.131 | −0.212 | 0.305 |
| | mass | −0.017 | 0.080 | −0.175 | 0.138 |
| | **forearm** | **0.266** | **0.150** | **0.006** | **0.589** |
| *silvivagum* | **$Bat_{castanea}$** | **−0.336** | **0.130** | **−0.582** | **−0.068** |
| | **$Bat_{perspicillata}$** | **0.446** | **0.123** | **0.200** | **0.680** |
| | $Bat_{sowelli}$ | −0.110 | 0.110 | −0.324 | 0.106 |
| | $Sex_{male}$ | 0.057 | 0.076 | −0.091 | 0.205 |
| | $Age_{juvenile}$ | −0.101 | 0.198 | −0.484 | 0.285 |
| | $Age_{subadult}$ | 0.014 | 0.211 | −0.405 | 0.424 |
| | $Rep_{reproductive}$ | 0.087 | 0.147 | −0.196 | 0.378 |
| | mass | −0.082 | 0.089 | −0.256 | 0.093 |
| | forearm | −0.068 | 0.097 | −0.249 | 0.133 |
| *umbricola* | **$Bat_{castanea}$** | **−0.182** | **0.133** | **−0.442** | **0.079** |
| | **$Bat_{perspicillata}$** | **0.197** | **0.125** | **−0.051** | **0.441** |
| | $Bat_{sowelli}$ | −0.015 | 0.100 | −0.213 | 0.181 |
| | $Sex_{male}$ | 0.121 | 0.072 | −0.019 | 0.263 |
| | $Age_{juvenile}$ | 0.108 | 0.173 | −0.231 | 0.449 |

| | | | | | |
|---|---|---|---|---|---|
| | $Age_{subadult}$ | 0.019 | 0.189 | −0.356 | 0.386 |
| | $Rep_{reproductive}$ | −0.127 | 0.125 | −0.373 | 0.120 |
| | mass | −0.133 | 0.080 | −0.293 | 0.025 |
| | forearm | 0.030 | 0.101 | −0.161 | 0.236 |
| *urostachyum* | $Bat_{castanea}$ | −0.164 | 0.130 | −0.417 | 0.090 |
| | $Bat_{perspicillata}$ | 0.219 | 0.122 | −0.019 | 0.456 |
| | $Bat_{sowelli}$ | −0.055 | 0.098 | −0.247 | 0.134 |
| | $Sex_{male}$ | 0.040 | 0.070 | −0.096 | 0.177 |
| | $Age_{juvenile}$ | −0.024 | 0.177 | −0.370 | 0.323 |
| | $Age_{subadult}$ | −0.009 | 0.195 | −0.381 | 0.376 |
| | $Rep_{reproductive}$ | 0.032 | 0.132 | −0.222 | 0.292 |
| | mass | −0.061 | 0.080 | −0.217 | 0.094 |
| | forearm | 0.123 | 0.097 | −0.067 | 0.313 |
| Type 1 | $Bat_{castanea}$ | 0.251 | 0.156 | −0.058 | 0.549 |
| | **$Bat_{perspicillata}$** | **−0.292** | **0.145** | **−0.576** | **−0.007** |
| | $Bat_{sowelli}$ | 0.041 | 0.097 | −0.150 | 0.229 |
| | $Sex_{male}$ | −0.072 | 0.070 | −0.208 | 0.067 |
| | $Age_{juvenile}$ | 0.175 | 0.180 | −0.172 | 0.529 |
| | $Age_{subadult}$ | 0.034 | 0.199 | −0.361 | 0.416 |
| | $Rep_{reproductive}$ | −0.210 | 0.131 | −0.462 | 0.048 |
| | mass | 0.081 | 0.079 | −0.076 | 0.234 |
| | **forearm** | **0.699** | **0.176** | **0.341** | **1.016** |
| Type 4 | **$Bat_{castanea}$** | **−0.318** | **0.119** | **−0.540** | **−0.080** |
| | **$Bat_{perspicillata}$** | **0.361** | **0.112** | **0.132** | **0.561** |
| | $Bat_{sowelli}$ | −0.043 | 0.094 | −0.228 | 0.142 |
| | $Sex_{male}$ | −0.015 | 0.066 | −0.145 | 0.114 |
| | $Age_{juvenile}$ | −0.103 | 0.176 | −0.445 | 0.239 |
| | $Age_{subadult}$ | 0.017 | 0.193 | −0.357 | 0.400 |
| | $Rep_{reproductive}$ | 0.085 | 0.127 | −0.160 | 0.338 |
| | mass | −0.013 | 0.075 | −0.162 | 0.135 |
| | **forearm** | **−0.213** | **0.079** | **−0.366** | **−0.056** |
| Type 8 | $Bat_{castanea}$ | −0.234 | 0.126 | −0.474 | 0.021 |
| | $Bat_{perspicillata}$ | 0.225 | 0.122 | −0.017 | 0.463 |
| | $Bat_{sowelli}$ | 0.010 | 0.098 | −0.181 | 0.203 |
| | $Sex_{male}$ | 0.040 | 0.070 | −0.099 | 0.173 |
| | $Age_{juvenile}$ | −0.006 | 0.184 | −0.371 | 0.353 |
| | $Age_{subadult}$ | 0.025 | 0.200 | −0.366 | 0.419 |

| | | | | | |
|---|---|---|---|---|---|
| | $Rep_{reproductive}$ | −0.019 | 0.133 | −0.282 | 0.239 |
| | mass | −0.026 | 0.080 | −0.180 | 0.131 |
| | forearm | 0.102 | 0.098 | −0.089 | 0.295 |
| Type 10 | $Bat_{castanea}$ | −0.269 | 0.169 | −0.587 | 0.069 |
| | **$Bat_{perspicillata}$** | **0.307** | **0.149** | **0.001** | **0.590** |
| | $Bat_{sowelli}$ | −0.037 | 0.110 | −0.251 | 0.180 |
| | $Sex_{male}$ | −0.035 | 0.077 | −0.185 | 0.116 |
| | $Age_{juvenile}$ | 0.048 | 0.193 | −0.334 | 0.421 |
| | $Age_{subadult}$ | −0.129 | 0.210 | −0.537 | 0.284 |
| | $Rep_{reproductive}$ | 0.081 | 0.142 | −0.199 | 0.359 |
| | **mass** | **−0.184** | **0.090** | **−0.364** | **−0.009** |
| | forearm | 0.231 | 0.172 | −0.068 | 0.602 |
| Type 11 | $Bat_{castanea}$ | −0.011 | 0.168 | −0.326 | 0.328 |
| | $Bat_{perspicillata}$ | 0.190 | 0.150 | −0.113 | 0.475 |
| | $Bat_{sowelli}$ | −0.179 | 0.111 | −0.398 | 0.037 |
| | $Sex_{male}$ | 0.110 | 0.079 | −0.043 | 0.263 |
| | $Age_{juvenile}$ | −0.063 | 0.199 | −0.453 | 0.330 |
| | $Age_{subadult}$ | 0.016 | 0.211 | −0.402 | 0.434 |
| | $Rep_{reproductive}$ | 0.047 | 0.144 | −0.237 | 0.332 |
| | mass | −0.069 | 0.091 | −0.250 | 0.107 |
| | forearm | 0.291 | 0.180 | −0.028 | 0.667 |

**Table S3**. Results from hierarchical maximum likelihood regressions of bite force as a function of bat individual and species traits. Each covariate corresponds to a mean measurement per bat species *i* in species *k*. Deviance is a measure of how well the model fits the data, with higher values of deviance indicating the data deviates substantially from model predictions. Estimates in bold a 95% posterior distribution that does not cross zero.

| Formula | Estimate | Mean | 2.50% | 97.50% |
|---|---|---|---|---|
| $bite.force_i \sim \alpha_k + \beta_1 * head.length_i + \beta_2 * body.mass_i + \beta_3 * male.sex_i + \varepsilon_i + \varepsilon_k$ | | | | |
| | $\boldsymbol{\alpha_{castanea}}$ | **−4.686** | **−9.345** | **−0.273** |
| | $\alpha_{perspicillata}$ | −3.933 | −8.828 | 0.639 |
| | $\alpha_{sowelli}$ | −3.899 | −8.737 | 0.625 |
| | $\boldsymbol{\beta_1}$ | **1.932** | **0.434** | **3.517** |
| | $\beta_2$ | −0.074 | −0.51 | 0.341 |
| | $\boldsymbol{\beta_3}$ | **0.412** | **0.194** | **0.623** |
| | $\varepsilon_i$ | 0.162 | 0.118 | 0.229 |
| | $\varepsilon_k$ | 2.623 | 0.259 | 16.003 |
| | Deviance | −21.808 | −28.401 | −9.991 |
| $bite.force_i \sim \alpha_k + \beta_1 * head.length_i + \beta_2 * forearm.length_i + \beta_3 * male.sex_i + \varepsilon_i + \varepsilon_k$ | | | | |
| | $\alpha_{castanea}$ | 3.151 | −5.756 | 11.161 |
| | $\alpha_{perspicillata}$ | 4.165 | −5.165 | 12.597 |
| | $\alpha_{sowelli}$ | 4.024 | −5.195 | 12.307 |
| | $\boldsymbol{\beta_1}$ | **1.470** | **0.281** | **2.718** |
| | $\beta_2$ | −1.772 | −4.15 | 0.669 |
| | $\boldsymbol{\beta_3}$ | **0.270** | **0.140** | **0.403** |
| | $\varepsilon_i$ | 0.155 | 0.117 | 0.21 |
| | $\varepsilon_k$ | 2.971 | 0.306 | 18.582 |
| | Deviance | −28.504 | −34.851 | −17.332 |
| $bite.force_i \sim \alpha_k + \beta_1 * head.length_i + \beta_3 * male.sex_i + \varepsilon_i + \varepsilon_k$ | | | | |
| | $\alpha_{castanea}$ | −2.34 | −5.828 | 1.251 |
| | $\alpha_{perspicillata}$ | −1.599 | −5.266 | 2.199 |
| | $\alpha_{sowelli}$ | −1.63 | −5.26 | 2.074 |
| | $\boldsymbol{\beta_1}$ | **1.176** | **0.005** | **2.328** |
| | $\boldsymbol{\beta_3}$ | **0.261** | **0.127** | **0.401** |
| | $\varepsilon_i$ | 0.16 | 0.12 | 0.215 |
| | $\varepsilon_k$ | 3.258 | 0.249 | 19.544 |

| | | | |
|---|---|---|---|
| Deviance | −26.424 | −31.925 | −16.462 |

**Table S4.** Results from phylogenetic Bayesian regressions of GJAM *Piper* consumption indices for *C. castanea* as a function of *Piper* species traits. Each covariate corresponds to a mean measurement per *Piper* species *i*. $Piper^{Cca}$ represents the proclivity of *castanea* for each *Piper* species estimated by the GJAM model. Each covariate corresponds to a mean by *Piper* species *i*. $\alpha$, intercept, $\beta$ coefficient, $\Sigma$ phylogenetic variance, $\varepsilon_i$ residual variance.

| formula | parameter | mean | 2.5% | 97.5% |
|---|---|---|---|---|
| $Piper^{Cca}{}_i \sim \alpha + \beta *$ seed ratio$_i + \Sigma + \varepsilon_i$ | intercept ($\alpha$) | −0.083 | −0.724 | 0.719 |
| | $\beta$ | −0.052 | −0.572 | 0.457 |
| | $\Sigma$ | 0.039 | 0.000 | 0.160 |
| | $\varepsilon$ | 0.160 | 0. 051 | 0.317 |
| $Piper^{Cca}{}_i \sim \alpha + \beta *$ infructescence.ratio$_i + \Sigma + \varepsilon_i$ | intercept ($\alpha$) | −0.045 | −0.491 | 0.458 |
| | $\beta$ | −0.016 | −0.016 | 0.017 |
| | $\Sigma$ | 0.028 | 0.000 | 0.122 |
| | $\varepsilon$ | 0.131 | 0.038 | 0.268 |

**Figure S1.** Illustration of A) lateral view of *Carollia castanea* head cranium and dorsal view showing linear measurements of head dimensions taken from live bats. HL is head length, HH is head height, HW is head width and B) set up for measuring bite force from bats *C. castanea.* Photo credit: David Villalobos Chavez.

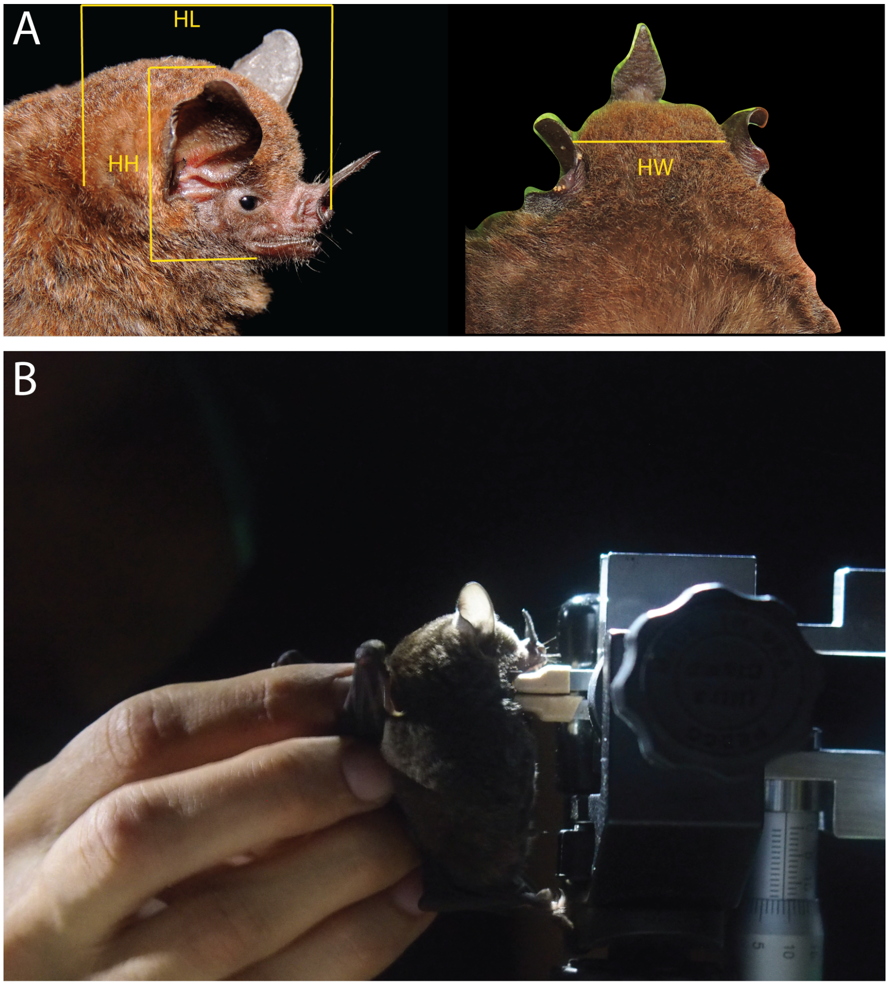

**Figure S2.** Percentage of non–*Piper* and *Piper* species in the diet of three *Carollia* bats. Note that Type 7, *P. auritifolium*, *P. biolleyi*, *P. decurrens*, *P. peltatum*, *P. terrabanum*, and *P. trigonum* have been removed as they account for less than 1% of the diet. See Data S1 for full spreadsheet.

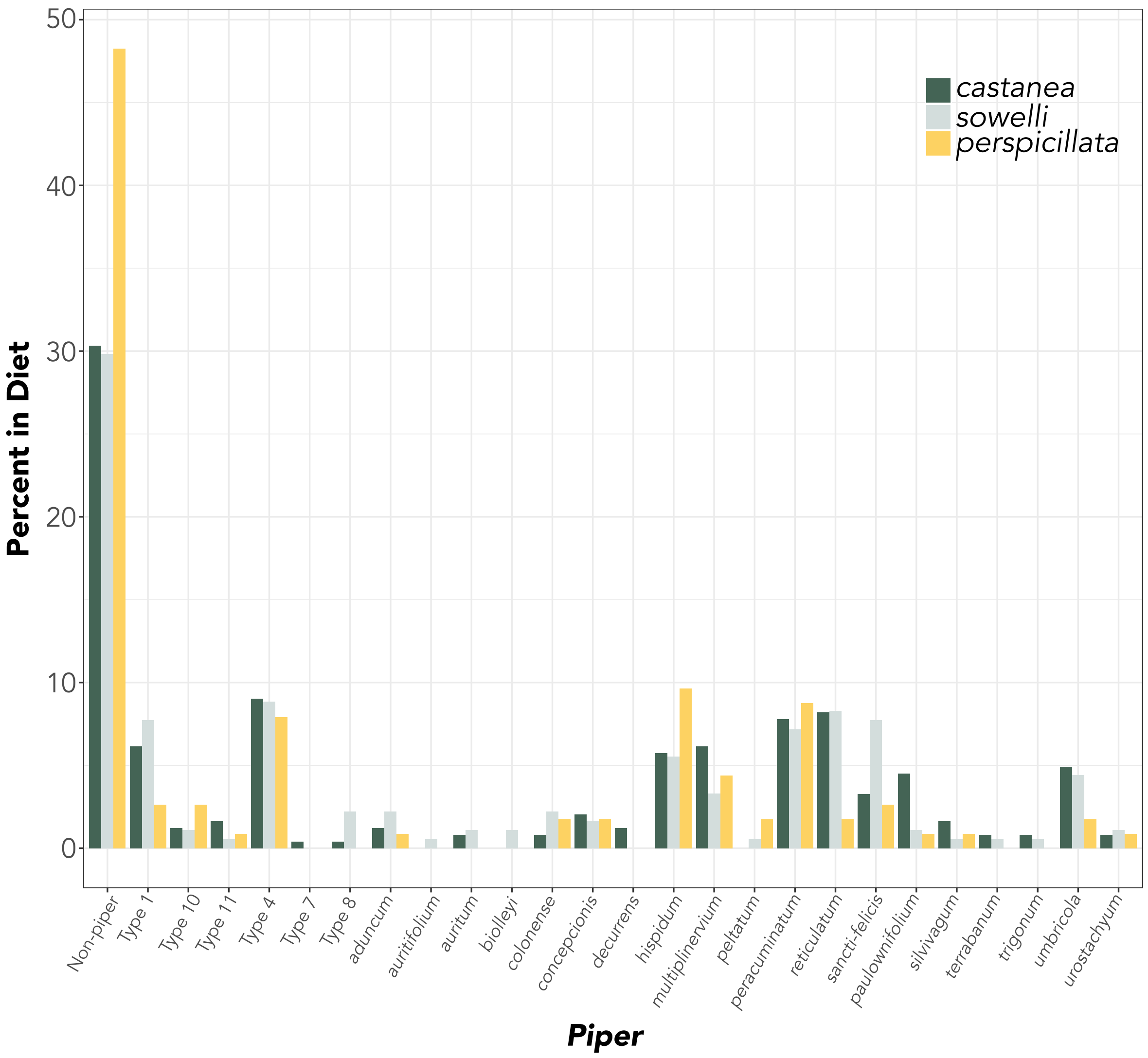

**Figure S3**. Body size measurements for three bat species, showing the variation in body mass (A) and forearm length (B). Forearm length is a standard and often diagnostic measure of size for bats.

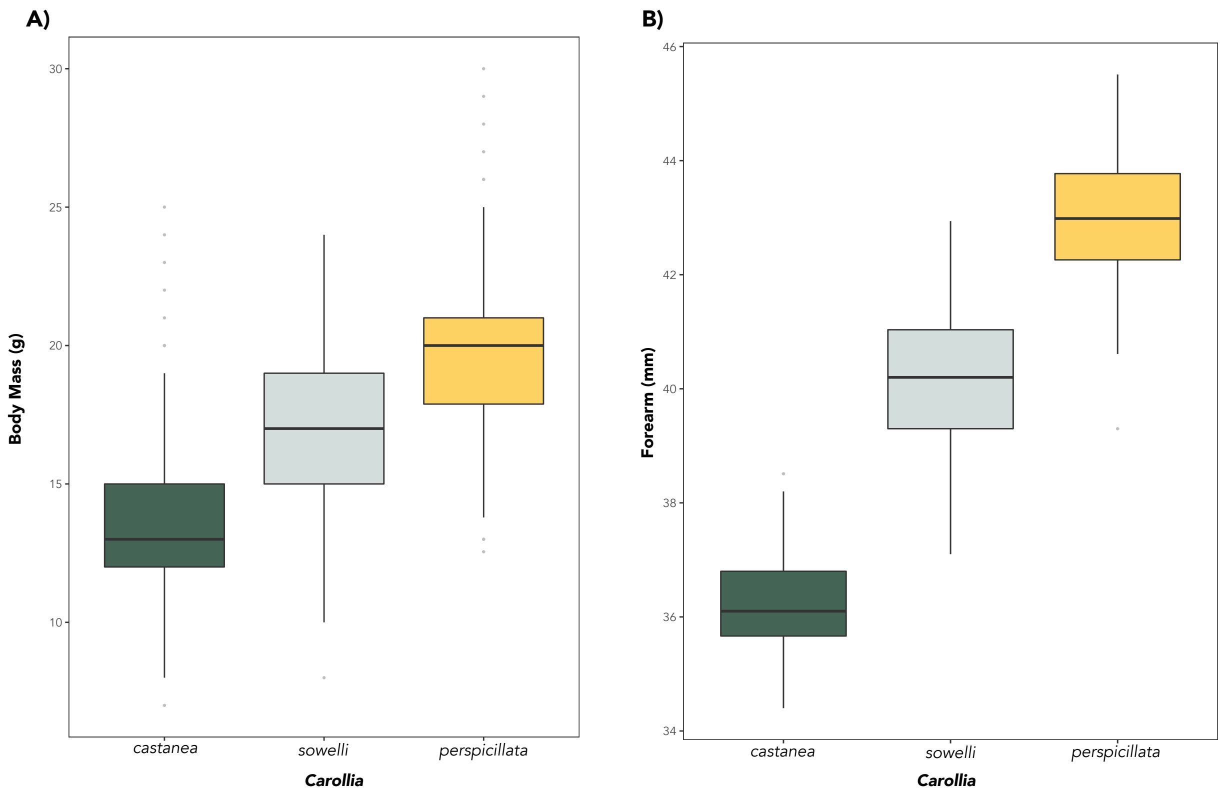

**Figure S4**. Posterior distributions of age responses for each *Piper* species, ordered by median. Probabilities represent the influence of age category on *Piper* consumption index.

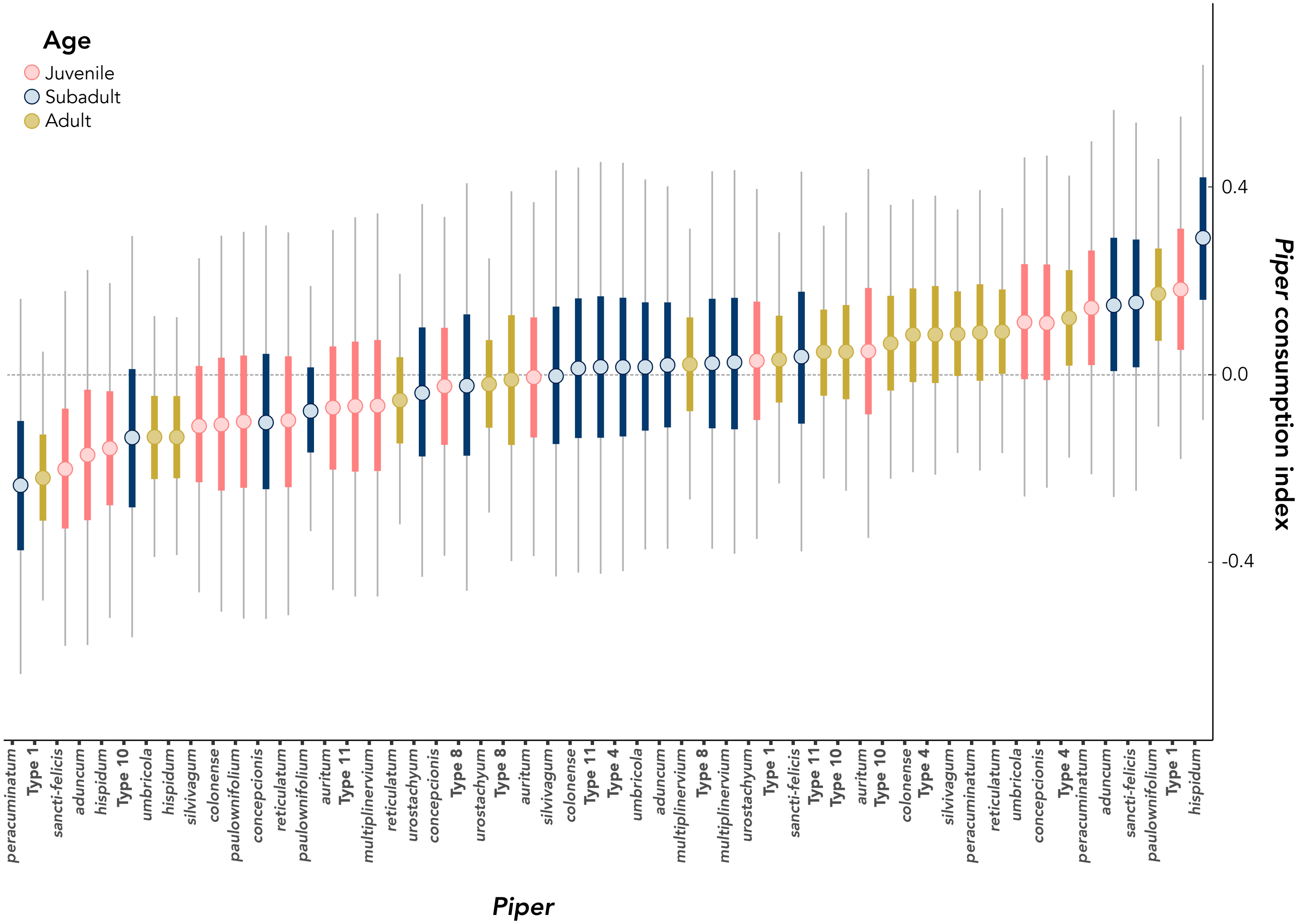
